# Distracting stimuli evoke ventral tegmental area responses in rats during ongoing saccharin consumption

**DOI:** 10.1101/2020.08.03.228452

**Authors:** Kate Z Peters, Andrew M J Young, James E McCutcheon

## Abstract

Disruptions in attention, salience and increased distractibility are implicated in multiple psychiatric conditions. The ventral tegmental area (VTA) is a potential site for converging information about external stimuli and internal states to be integrated and guide adaptive behaviours. Given the dual role of dopamine signals in both driving ongoing behaviours (e.g. feeding) and monitoring salient environmental stimuli, understanding the interaction between these functions is crucial. Here we investigate VTA neuronal activity during distraction from ongoing feeding. We developed a task to assess distraction exploiting self-paced licking in rats. Rats trained to lick for saccharin were given a distraction test, in which three consecutive licks within 1 second triggered a random distractor (e.g. light and tone stimulus). On each trial they were quantified as distracted or not based on the length of their pauses in licking behaviour. We expressed GCaMP6s in VTA neurons and used fibre photometry to record calcium fluctuations during this task as a proxy for neuronal activity. Distractor stimuli caused rats to interrupt their consumption of saccharin, a behavioural effect which quickly habituated with repeat testing. VTA neural activity showed consistent increases to distractor presentations and, furthermore, these responses were greater on distracted trials compared to non-distracted trials. Interestingly, neural responses show a slower habituation than behaviour with consistent VTA responses seen to distractors even after they are no longer distracting. These data highlight the complex role of the VTA in maintaining ongoing appetitive and consummatory behaviours while also monitoring the environment for salient stimuli.

## Introduction

To operate effectively in an unpredictable world it is essential for animals to learn to pursue rewards and avoid potential threats and punishments. This requires both moment-to-moment monitoring of environmental stimuli and longer-term learning and memory processes. In this framework, it is critical that a balance exists between continuing to engage in rewarding behaviour (such as maintaining ongoing behaviour such as feeding) whilst also maintaining vigilance (monitoring the environment for potential new opportunities or threats).

This exploitation/exploration conflict is a vital part of adaptive behaviour in animals and humans. Deciding when to switch current behavioural output towards other stimuli in a changing environment is a central tenet of behavioural flexibility (Haluk & Floresco, 2009; Floresco, 2013). Adapting to changing environments is crucial to ensure that behaviours such as feeding are efficient and immediate survival needs are met but without leaving an animal vulnerable to danger such as attack from a predator. To achieve this, animals must be equipped with neural systems to both invigorate and sustain behaviours resulting in positive outcomes whilst at the same time supressing those that result in punishment (Cools *et al*., 2011). Classically, it is the dopamine system that has been implicated in behavioural activation (Salamone *et al*., 2009) and reward (Wise, 2004; Schultz, 2007, 2017; Flagel *et al*., 2011). When learning about rewards, not only is dopamine responsible for invigorating behavioural output but the precise timing of dopamine neuron firing encodes a ‘reward prediction error’ (Schultz, 1997, 2002). Reward prediction error acts as a teaching signal to compare expected and acquired rewards to promote learning and maximise reward. A related view of dopamine describes it as the ‘flexibility transmitter’ and is involved in behavioural switching (Beeler *et al*., 2014).

In uncertain environments, dopamine not only signals reward predictions but also encodes novel cues which are seemingly neutral (not associated with any type of reinforcer) (Ljungberg *et al*., 1992; Horvitz *et al*., 1997). In this case, short latency dopamine bursts (Redgrave *et al*., 1999) may signal the potential importance or salience of stimuli to guide behaviour towards the most reinforcing actions (Kakade & Dayan, 2002; Lisman & Grace, 2005). Midbrain dopamine encoding of novelty is important for learning (Waelti *et al*., 2001) and optogenetic activation of dopamine neurons reproduces the effects of novel stimuli, signalling new information and promoting reward learning (Steinberg *et al*., 2013). Such responses to novel stimuli might favour exploration rather than exploitation of existing resources when new information is available (Kakade & Dayan, 2002; Bromberg-Martin & Hikosaka, 2009). Midbrain dopamine cell firing can encode an initial salience (intensity and importance) and surprise (the presence of new or unexpected stimuli) prior to these stimuli becoming associated with reward/value (Nomoto *et al*., 2010; Fiorillo *et al*., 2013). The dopamine system in particular seems to encode multiple aspects of reward environments across time, from salience to motivation to learned associations (Bromberg-Martin, Matsumoto, & Hikosaka, 2010; Bromberg-Martin, Matsumoto, Nakahara, *et al*., 2010). The VTA in particular is a potential hub for the integration of these multiplex signals, where inputs from cortical and other areas (such as the serotonergic and glutamate neurons of the dorsal raphe nucleus (Liu *et al*., 2014; Wang *et al*., 2019) may converge to modulate attention and adaptive decision making.

The processes of attention and salience are key to the balance between exploiting current opportunities and exploring others. When appropriately deployed, attention to salient stimuli allows an animal to focus on a certain task while successfully monitoring the surrounding world to rapidly switch behaviours if necessary. Exactly how the sensory aspects of stimuli are assimilated into ongoing behaviour through learned associations is still not fully understood. While reacting to stimuli in the outside world at appropriate times will act to protect an animal from danger, over-vigilance and distraction by external stimuli is maladaptive and will lead to an animal failing to exploit environmental resources effectively (Riccio *et al*., 2002; Moher *et al*., 2015).

Finally, it is important to recognise that plasticity and learning in this system is critical. As such, a stimulus that might capture attention on its first presentation should, over repeated presentations become less salient to an animal and thus become less likely to disrupt ongoing behaviour. Promoting flexible approach behaviours involves rapid assimilation of external stimuli, at times from multiple modalities, and decision making based on probabilities of opportunity, calculations of cost and effort as well as balancing ongoing behaviours with possible future rewards from new opportunities. Moreover, traits such as hyper-vigilance, inappropriate allocation of attention, disrupted novelty detection and increased distraction in humans are hallmarks of several psychiatric diseases including schizophrenia, post-traumatic stress disorder, addiction and anxiety disorders (Winton-Brown *et al*., 2014).

Here, we have developed a behavioural paradigm in which we monitor rats’ ongoing consumption of a palatable solution while presenting external, distracting stimuli that are triggered by their licking behaviour. In the same rats, we recorded neural signals from VTA using fibre photometry. We aimed to investigate how external distractors affect ongoing consumption and neural activity. Our hypotheses are that presentation of external stimuli will lead to interruptions in ongoing licking behaviour but that these interruptions will habituate over time. Furthermore, we predict that neural activity in VTA will reflect aspects of the ongoing licking behaviour as well as responding to distracting stimuli and, importantly, that whether an animal is distracted or not will be reflected in the neural activity evoked by the stimulus.

## Materials and methods

### Animals

Thirteen adult male Sprague Dawley rats were used in behavioural and photometry experiments (Charles River, UK: 300g - 350g at time of surgery). One rat was removed from analysis because it was not possible to make fibre photometry recordings on all test days. All rats were housed in pairs in individually ventilated cages under temperature controlled conditions (21 °C ± 2°C; 40-50% humidity) and kept under 12 h light/dark cycle, with lights on at 07:00. Post-surgery, rats were housed with bedding materials recommended by the NC3Rs and never single housed. Chow and water were available ad libitum except for a brief period of food restriction (12 hours before the first training session) and during experimental sessions when only saccharin solution was given (1 h per day). All procedures were carried out under the appropriate UK government Home Office license authority (PPL #70/8069) in accordance with the Animals [Scientific Procedures] Act (1986).

### Virus injection and implant surgery

Rats were deeply anaesthetized using isoflurane (5% / 2 L/min for induction, 2% for maintenance). The head was shaved and pre-operative analgesia was administered (bupivacaine 150 μL at incision site and meloxicam 1 mg/kg s.c.). Rats were mounted in a stereotaxic frame (David Kopf Instruments) and a thermostatic blanket was used to maintain a stable body temperature throughout surgery (37-38 °C). Constant monitoring of oxygen saturation and heart rate was performed with a pulse oximeter. A scalp incision was made and a hole was drilled above the VTA for virus injection and fibre implantation (from Bregma: AP: −5.8, ML: +0.7). In addition, holes were drilled anterior and posterior to attach four anchor screws. A virus injection needle (10 μl Hamilton Syringe) was lowered into the VTA (DV −8.1 mm, from dura) and 1 μl of virus (AAV9.Syn.GCaMP6s.WPRE.SV40, ~1.9×10^13^ GC/ml; Penn Vector Core) was injected over 10 min using a syringe pump (11 Elite, Harvard Apparatus, CA). A fibre optic cannula was implanted at DV −8.0 mm (0.1 mm above the virus injection site) (ThorLabs CFM14L10, 400 μm, 0.39 NA, 10 mm, sterilized using ethylene oxide). A layer of radio-opaque dental cement was used to seal screws and cannula in place (C&B Super-Bond, Prestige Dental) and a headcap was formed using dental acrylic (DuraLay, Reliance Dental). Care was taken to leave approximately 5 mm of the ferrule protruding for coupling to optical patch cable for later photometry recordings. Rats were housed in pairs immediately following surgery and at least 4 weeks was allowed post-surgery for virus expression before testing began.

### Fibre photometry

Fibre photometry equipment was similar to that reported elsewhere (Lerner *et al*., 2015) and consisted of two fibre coupled light sources powered by LED drivers. A blue 470 nm LED (Thorlabs, M470F3) and violet 405 nm LED (Thorlabs, M405F1) were sinusoidally modulated at 211 Hz and 539 Hz, respectively, and passed through filters (470 nm and 405 nm). Both light paths were directed, via dichroic mirrors positioned inside filter cubes (FMC4_AE(405)_E(460-490)_F(500-550)_S, Doric Lenses), through a fibre optic patch cord (MFP_400/460/LWMJ-0.48_3.5m_FCM_MF2.5, Doric Lenses). The patch cord was then mated, using a ceramic sleeve, to the implanted fibre optic cannula. Emitted fluorescence was collected via the same fibre, through the patch cord and focused onto a photoreceiver (2151, Newport). A signal processor (RZ5P; Tucker Davis Technologies) and Synapse software (Tucker Davis Technologies) was used to control LEDs, to acquire the lowpass filtered signal (3 Hz), and to perform on-line demodulation of the signal, separating isosbestic and calcium-modulated responses. Demodulated signals were acquired at 1017 Hz. Behavioural events, such as licks and distractors, were routed to this system as TTLs and acquired simultaneously.

### Distraction testing

Behavioural experiments were carried out in operant behaviour chambers (Med Associates, VT, USA; 25 cm X 32 cm X 25.5 cm) housed inside large sound attenuating chambers. On one wall of the chamber there was a sipper with a cue light above it. A grid floor, comprised of stainless steel rods, was used in conjunction with contact lickometers, to record individual licks as rats consumed solutions from a spout recessed 5-10 mm from the chamber wall. A circular hole in the ceiling of the operant chamber allowed the patch cord to pass through allowing full, free movement of the animal during photometry recording sessions. Rats were initially trained to lick for sodium saccharin solution (0.2% in distilled water; Sigma 47839) during 60 min sessions, with saccharin freely available from the spout. To encourage rats to start licking they were food restricted overnight, 12 h prior to the first licking session only. Rats were trained for 3-6 days until they reached a criterion of 1000 licks in a single session.

Distraction test days were identical to lick training days, except the house light was not illuminated and hardware was configured to deliver distracting stimuli (distractors). Distractors were triggered following 3-5 consecutive licks within 1 s (Fig. 1A). Distractors lasted for 1 s and were pseudorandomly chosen from the following list of stimuli: a cue light, flashing cue light, tone (5 kHz, 80 dB), burst of white noise (flat 10 – 20 kHz) or combination of these. All distractors contained both an audible and a visual stimulus. If rats paused licking following the distractor for >1 s they were deemed to be distracted and, if not, they were deemed not distracted on that trial. To avoid rats receiving distractors too frequently, i.e. during long trains of licks, rats had to pause licking for over 1 s before the next distractor was presented. All distractors were found to be effective at causing distraction according to this definition on a proportion of trials, however, distractors that included white noise were found to be marginally more effective than those that just included the tone (Fig. S1; F(2,24)=4.40, p=0.0235). No difference was found on their ability to modulate neural activity (F(2,24)=0.28, p=0.7554). Rats were tested on the distraction test for two days to assess habituation to the task.

**Figure 1.**
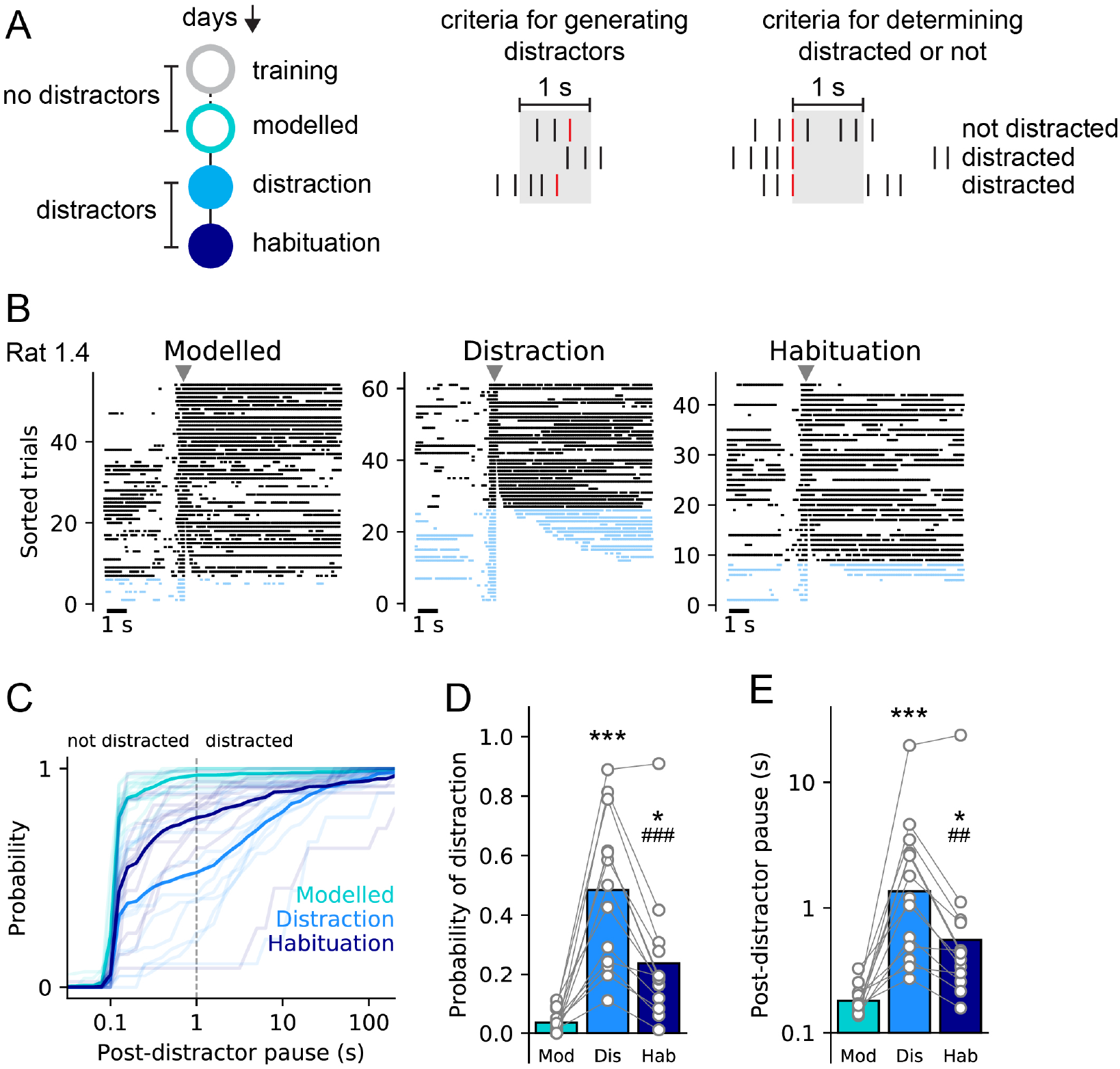
Licking behaviour differs between days when distractor is presented and when it is not. (A) Schematics showing experimental timeline (left), criteria for generating distractors (centre), and criteria for determining whether presentation of a distractor caused distraction or not (right). Licks are shown as vertical lines with red lines indicating licks that trigger a distractor. (B) Representative raster plots showing licks aligned to distractor and sorted by the post-distractor pause. On modelled day, no distractors were presented but time when distractors would have occurred was calculated. Not distracted trials are shown in black and distracted trials are shown in blue. Grey triangles show time of distractor. (C) Cumulative probability of post-distractor pause durations across days. Solid lines show mean of all rats and light lines show individual rats. Dashed vertical line denotes criteria (1 s) for determining trials on which a rat was classified as distracted or not. (D) Probability of distraction is greater on first distraction day than on modelled day or habituation day. (E) Post-distractor pause is longer on first distraction day than on modelled day or habituation day. For D and E circles show data from individual rats and bars are mean. ***, * p<0.001, p<0.05 vs. modelled day; ###, ## p<0.001, p<0.01 vs. distraction day.

### Modelled distractors

To confirm that the distraction task did not simply capture a normal pattern of licking (i.e., to verify that animals do not – independently of distractors – show patterns of 3-5 lick bursts with pauses), we programmatically determined when rats would have received distractors during normal licking sessions. This allowed us to determine, according to the interlick interval following the distractor, if rats would have been classified as distracted or not and thus provided a baseline on lick training days against which we could compare behaviour from distraction days. Modelled distractors on lick days were compared to real distractors on distraction days.

### Histology

Immunohistochemistry was used to verify fibre implant sites in the VTA and to check the extent of regional viral expression. At the end of experiment, all rats were terminally anesthetised with isofluorane and sodium pentobarbital (5 ml/kg) before being transcardially perfused with ice-cold phosphate-buffered saline (PBS) followed by 4% paraformaldehyde (PFA). Following perfusion, brains were removed and placed in 4% PFA for 24 hours before being transferred to a cryoprotectant 30% sucrose solution in PBS for at least 3 days. 40 μm coronal sections were made using a freezing microtome and stored in 5% sucrose solution in PBS with 0.02% sodium azide until staining.

Free-floating sections were first incubated in blocking solution (3% normal goat serum, 3% normal donkey serum, 3% Triton-X in PBS) for 1 h before incubation with primary antibodies (1:1000 anti-tyrosine hydroxylase, #AB152, Millipore UK; 1:1000 anti-GFP, #A10262, Fisher Scientific, UK) in blocking solution at room temperature on a shaker for 18 h. Next, sections were incubated with secondary antibodies (1:250 donkey anti-Rabbit IgG 594, AlexaFluor, Fisher Scientific, UK and 1:250 goat anti-Chicken IgY 488, Life Science Technologies) in PBS for 90 min at room temperature. Sections were mounted on slides using Vector Shield hard-set mountant (Vector Labs, UK). Between all steps, sections were washed three times with PBS for 5 min and gently agitated on a laboratory shaker. Slices were imaged using an epifluorescent microscope (Leica, UK) to determine fibre placement and virus expression with the Paxinos & Watson (2007) rat brain atlas used as reference.

### Statistical analyses

All photometry and behavioural data were extracted from TDT files and analysed using custom Python scripts available at https://github.com/jaimemcc/Distraction-Paper/releases/tag/v2.0. These extracted data included signals for both photometry streams (calcium-modulated and isosbestic light levels) and timings of individual licks (onsets and offsets of contact with lickometer) and distractors (time stamps of delivered distraction stimuli). Raw TDT data files and extracted data files are available at https://www.doi.org/10.25392/leicester.data.12732734. The calcium-modulated signal was corrected for artifacts and bleaching using the isosbestic signal as a reference (Lerner *et al*., 2015). Once data were extracted, the corrected photometry signal was aligned to individual events (distractors or modelled distractors) and normalised via z-scoring to a 5 s baseline before the event of interest. For statistical analyses all data were expressed as means and Python or SPSS.24 (Chicago, USA) was used to perform mixed ANOVAs or t-tests, and receiver-operator characteristic (ROC) analysis, where appropriate. All assumptions of sphericity, homogeneity of variance and normality were satisfied unless otherwise stated. Alpha was set at p < .05, all significance tests were two-tailed, and Bonferroni or Sidak was used to correct for multiple comparisons.

## Results

### Rats pause ongoing saccharin consumption in response to external distractor stimuli

First, rats were trained to lick for saccharin solution over several 1 h sessions (3-6 days). Once rats had reached a stable amount of licking they experienced a distractor session in which a distracting stimulus was presented every time they made 3-5 licks within 1 s (see *Methods* for full details and Fig. 1A). This session was followed the next day by a second identical distraction session (termed habituation; Fig. 1A). To quantify changes in behaviour between the sessions with distractors and the licking session that preceded them, we calculated when distractors would have occurred in the first session (hereafter called *modelled distractors*) and examined the likelihood of rats pausing their licking by chance after 3-5 licks even with no distractor present. Our analysis showed that there was a low, but non-zero probability, of rats pausing licking (defined as >1s) after 3-5 licks in this session (0.035 ± 0.01). However, in the sessions in which distractors were present the probability of rats pausing after a distractor was 0.484 ± 0.069 and 0.238 ± 0.061 for the first and second sessions, respectively. This pattern is demonstrated by the representative raster plots of licking aligned to distractors (Fig. 1B). Indeed, statistical analysis with one-way repeated measures ANOVA revealed that during distraction sessions rats’ likelihood of pausing changed, relative to the previous session when no distractors were present (Fig. 1D; F(2,24)=28.5, p<0.001). As such, the likelihood of distraction changed across the sessions with there being a greater percentage of distracted trials on the first distraction day (Bonferroni corrected, p=0.0001) and the second distraction day (p=0.017), relative to the day without distractors. Moreover, both within-session and across session habituation to distractors was identified. As such, dividing the distractors into terciles showed that rats had a higher probability of distraction earlier in the distraction session than later on (Fig. S2; F(2,24)=5.01, p=0.0152). In addition, across session habituation to the distractors was also observed as there was a significantly lower probability of rats being distracted on the second distraction day, relative to the first (p=0.0004).

In addition, we examined how duration of the pause after each distractor (or modelled distractor) changed across the three days (Fig. 1C and 1E). Analysis of log-transformed pauses revealed that the duration of this pause changed significantly across days (F(2,24)=23.0, p<0.0001). As such, the average pause was longer on both the first and second distraction days (p=0.0002 and p=0.017, respectively) than it was on the day without distracting stimuli. Again, demonstrating that habituation to the distractor occurred, the pause was shorter on the second distraction day than on the first distraction day (p=0.001).

We were also curious as to whether there were differences in licking behaviour – including in the period before the distracting stimulus - on trials in which rats were distracted vs. those on which they were not. To examine this, we pooled trials from the first distraction session for all rats and divided them based on our distraction criterion of a pause for >1 s following the distracting stimulus. We then conducted ROC analysis on binned licking data comparing distracted to non-distracted trials (Fig. 2A). As expected, based on our selection criteria, we found that there were significant differences between distracted and non-distracted trials in the 1 s following the distractor. In addition, there was no difference in the 2 s preceding distractor stimulus as criteria to trigger a distractor include a 1 s pause before a 3-5 lick burst. Interestingly, however, we found that these two classes of trials – distracted and non-distracted – also differed significantly both in the 15 s post-distractor period and in the 5 s pre-distractor period. In particular, differences in the period preceding the distractor indicate that the greater the rate of ongoing licking in the several seconds preceding occurrence of a distractor, the less likely a rat is to be distracted. We examined this across all rats by comparing the average lick rate before (Pre: −5 to 0 s) and after the distractor (Post: +1 to +15 s) between distracted and non-distracted trials (Fig. 2B). We found that the lick rate on distracted trials was significantly lower than on non-distracted trials both before (t(12)=2.6, p=0.021) and after presentation of the distractor (t(12)=7.6, p<0.0001).

**Figure 2.**
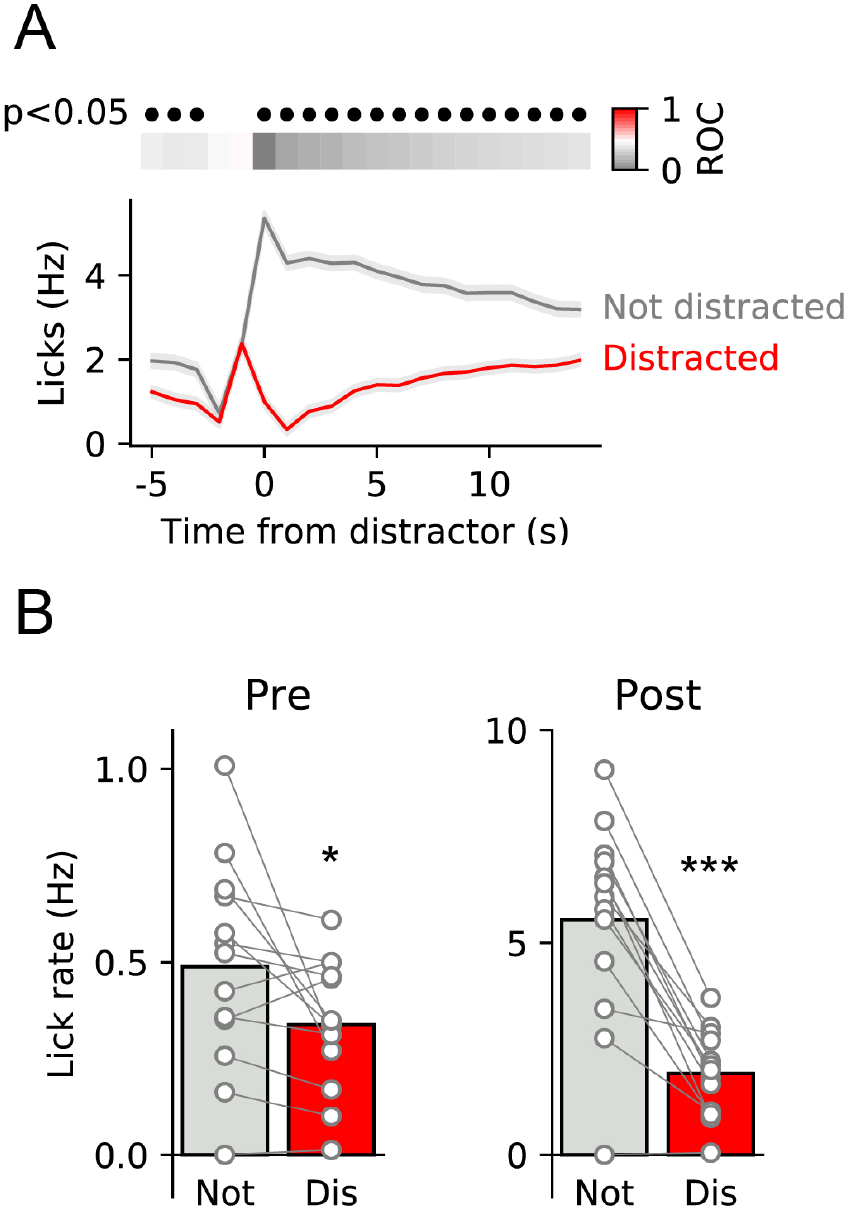
Licking behaviour on distracted trials differs from not distracted trials. (A) ROC analysis of distracted vs. not distracted trials shows that licking differs at all points except in the 2 s period when the distractor is triggered. Upper panel shows ROC values coded in colour with grey bins representing slower lick rates during distracted vs. not distracted trials. Black filled circles indicate time bins in which the ROC comparison was significant (Bonferroni corrected). Lower panel shows mean ± SEM of lick rates across all distracted and not distracted trials. (B) Comparisons of the mean lick rate before the distractor (left; Pre) and after the distractor (right; Post) for both types of trial show that lick rate was greater on not distracted trials for both these epochs. Circles show data from individual rats and bars are mean. *, *** p<0.05, p<0.001 vs. non-distracted trials.

The difference in lick rate before the distractor on distracted vs. non-distracted trials prompted us to explore this difference further. We considered whether the length of the pre-distractor pause was associated with whether the rat was distracted or not (Fig. S3). As might be predicted, we found that the pre-distractor pauses were shorter on non-distracted trials than on distracted trials (t-test of log-transformed data, p=2×10^-7^). As such, the median pre-distractor pause was 3.27 s on non-distracted trials vs. 5.57 s indicating that there was more likely to be a distractor occurring in this pre-distractor period on non-distracted trials than on distracted trials. In fact, on 64% of non-distracted trials there was a distractor occurring in the baseline period whereas for distracted trials this proportion was 41%.

In summary, our behavioural model shows that in many but not all cases presentation of a distracting stimulus will cause rats to pause in their ongoing licking and that this pause can vary greatly in its duration representing anything from a brief interruption to termination of a meal. The effectiveness of these distractors habituates relatively quickly so that in a second session, the distractors are far less likely to interrupt consumption. Finally, rats’ lick rate leading up to presentation of a distractor also seems to influence the effectiveness of a distracting stimulus with elevated lick rates being less likely to predict distraction

### VTA activity increases in response to distractor presentations

To assess how neural activity in VTA was associated with presentation of distracting stimuli we used fibre photometry to record fluctuations in calcium in VTA neurons during behaviour. Rats were injected with GCaMP6s in the VTA and fibre photometry cannulas were implanted 0.1 mm above virus injection (schematic in Fig. 3A). Histology confirmed that 13 probes were correctly placed and no animals were excluded due to incorrect placements or lack of viral expression at the fibre site (Fig. 3B shows probe locations).

**Figure 3.**
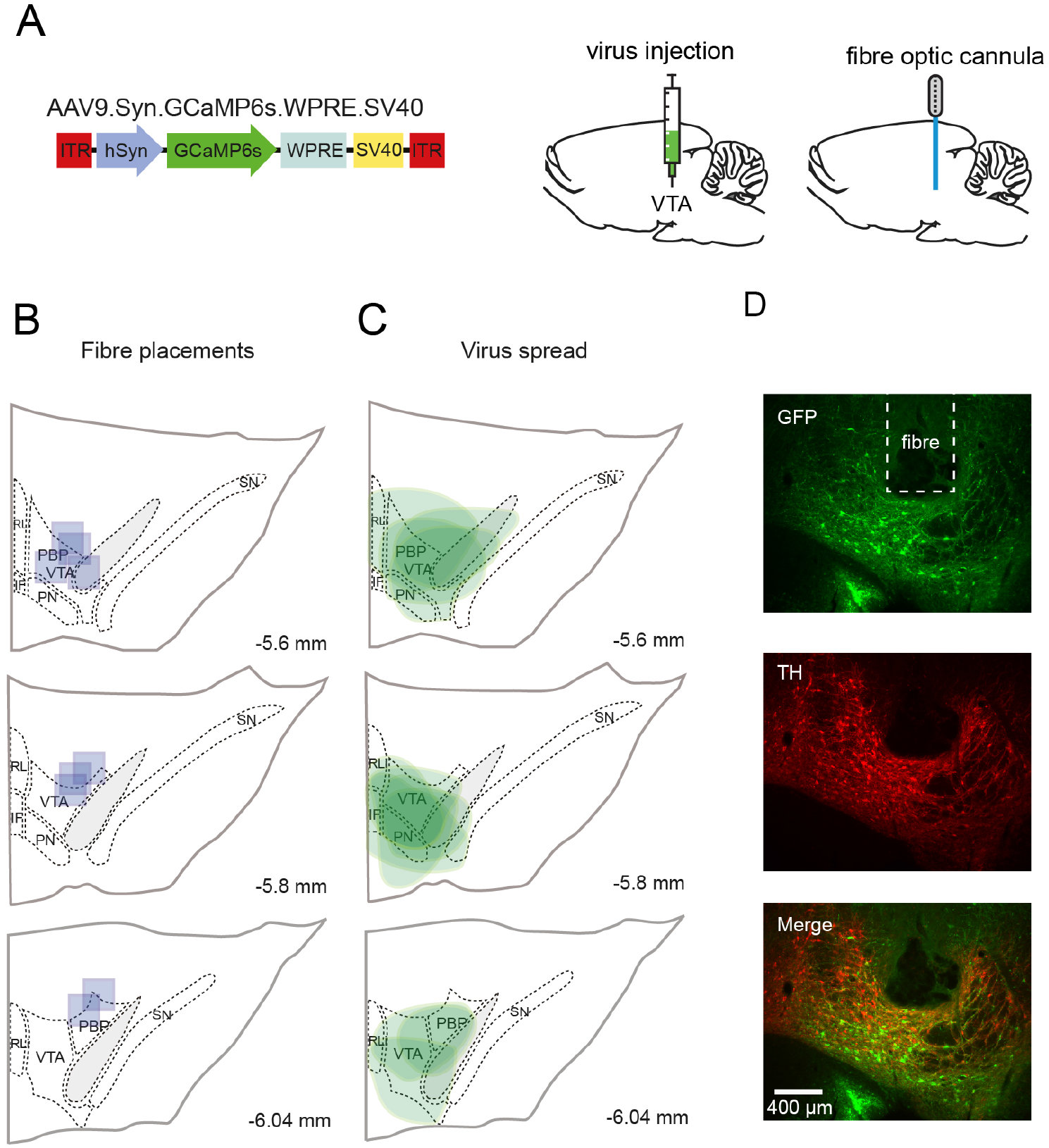
Viral injection and fibre placements within the VTA. (A) Schematic of GCaMP6s injections into the VTA of rats. Rats were injected with an adeno-associated virus carrying the calcium sensor and implanted in the VTA (0.1 mm above the viral injection site) with an optical fibre for photometry recordings. (B) Fibre placements within the VTA, each blue square shows the tip placement of the fibre optic for a single rat within the VTA, atlas images used for reference are adapted from Paxinos and Watson [2007] (C) Viral spread is highlighted in green, showing extent of GFP positive cells within the VTA. (D) Immunohistochemistry representative slice showing GFP expression (green), TH (red) and an overlay of both. Images taken at 10X magnification, inset square shows tip placement within that slice.

We first considered whether presentation of distracting stimuli affected neural activity by comparing the photometry signal between modelled day (when no distractors were present) and the first distraction day. For this comparison all photometry data were aligned to distractor or modelled distractor and converted to Z-scores. ROC analysis was used to compare each individual 1-s time bin from 5 s before until 15 s after each distractor on all trials from the modelled distractor and first distraction day (Fig. 4A). This analysis revealed that neural activity only differed in 3 consecutive time bins occurring following the distractor (Bonferroni corrected p’s<0.05). To confirm these findings, we examined this and the adjacent epochs by averaging selected bins and comparing the different trials across rats (Fig. 4B). As suggested by the ROC analysis, we found a significant interaction between epoch and day (two-way repeated-measures ANOVA: F(2,24)=10.96, p=0.0004). Specifically, in the epoch 1-4 s after the distractor neural activity was significantly elevated when distractors were present, relative to the modelled day that preceded it (Sidak: p=0.029). In contrast, in the period before the distractor (−5 to −1 s) and several seconds after the distractor (+5 to +15 s) there were no differences between the modelled and distraction days (p=0.962 and p=0.999, respectively). Thus, occurrence of distracting stimuli was associated with a transient (3 s) increase in neural activity.

**Figure 4.**
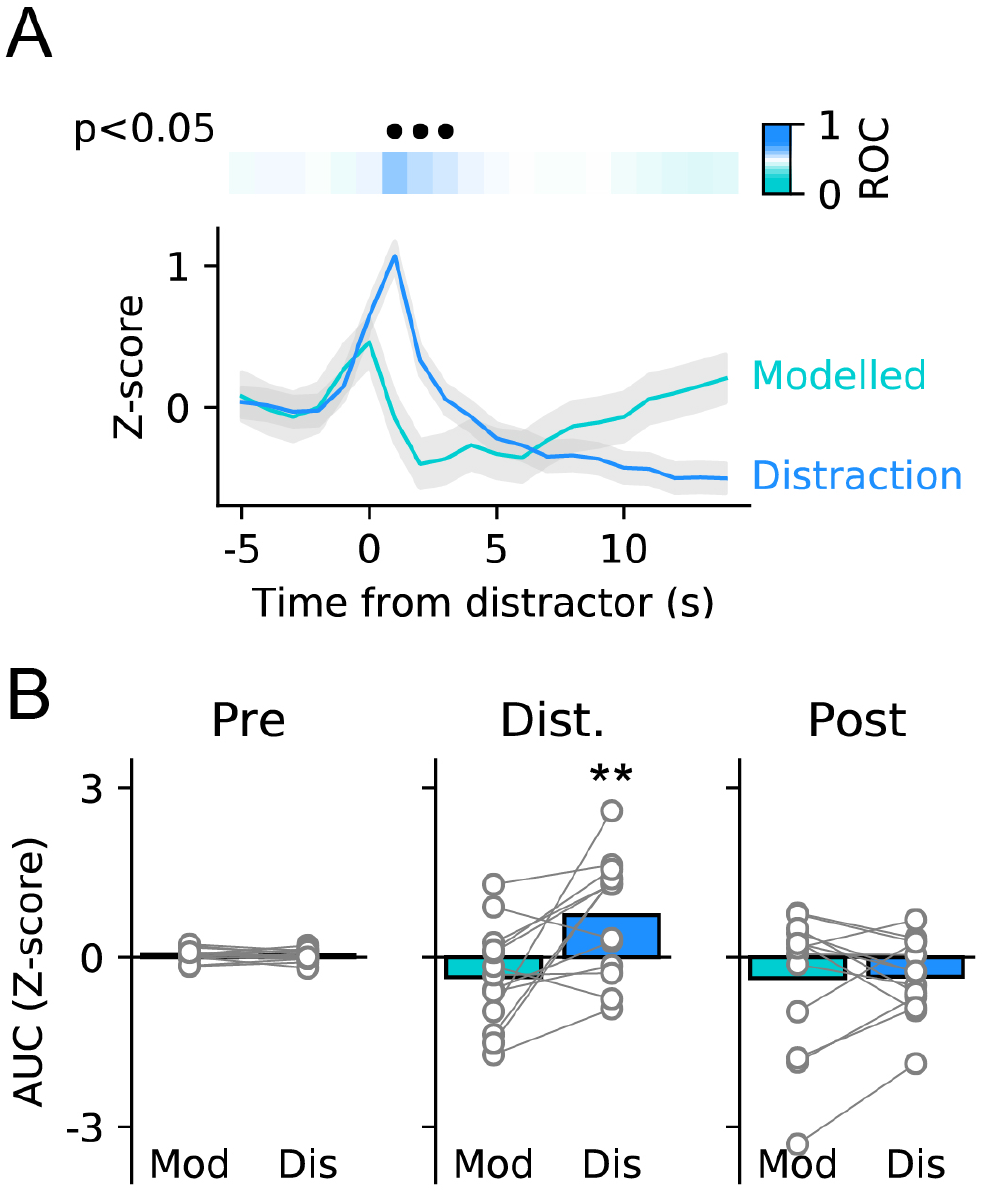
Neural activity is evoked by distracting stimuli. (A) ROC analysis of photometry data between modelled day with no distractors and first distraction day shows that neural activity differs in the 1.5 s following presentation of a distractor. Upper panel shows ROC values coded in colour with blue trials representing greater neural activity during real distractor trials vs. modelled distractor trials. Black filled circles indicate time bins in which the ROC comparison was significant (Bonferroni corrected). (B) Comparisons during three epochs show that neural activity is evoked in the epoch following the distractor (centre, 3 s following distractor) but does not differ before (left) or after (right) this period. Circles show data from individual rats and bars are mean. ** p<0.01 vs. modelled day.

### VTA responses are greater on distracted trials compared to non-distracted trials

Next, we wanted to know whether the neural response to distracting stimuli differed depending on whether rats were distracted (i.e. paused their licking) or not. To explore this we separated trials from the first distraction day into distracted and not distracted and performed ROC analysis on the aligned and normalised photometry signal (Fig. 5A). This analysis revealed that neural activity was significantly elevated for a prolonged time period following the distractor on distracted trials vs. non-distracted trials (Bonferroni corrected p’s<0.05). When we divided the trials into the same three epochs used before and compared across all rats, we found a significant interaction between epoch and trial type (two-way repeated-measures ANOVA: F(2,24)=10.85, p=0.0004). Comparisons of each epoch showed that there was no difference between activity in the baseline epoch (Sidak: 0.993), a trend towards a difference in the immediate epoch following the distractor (p=0.077), and a large difference in the later post-distractor epoch (p-0.0004), with activity being greater on distracted trials than non-distracted trials. As the rate of licking before the distractor was different in the 5 s period before presentation of the distractor, we also analysed the data before z-scoring (i.e. without normalisation to baseline) to examine whether there were pre-distractor differences in neural activity (Fig. S4). This analysis produced very similar results as our analysis of z-scored data (two-way repeated-measures, epoch x trial type interaction: F(2,24)=7.39, p=0.0032). Importantly, there was no difference in VTA activity before the distractor (Sidak: p=0.144) but differences in both post-distractor epochs (p=0.041 and p=0.007, respectively). Finally, for trials on which the rats were distracted, we tested whether the activity evoked by the distractor was related to the length of the pause before rats resumed licking. However, neural activity in the 3 s following distractors was not correlated with the length of the post-distraction pause (Fig. S5).

**Figure 5.**
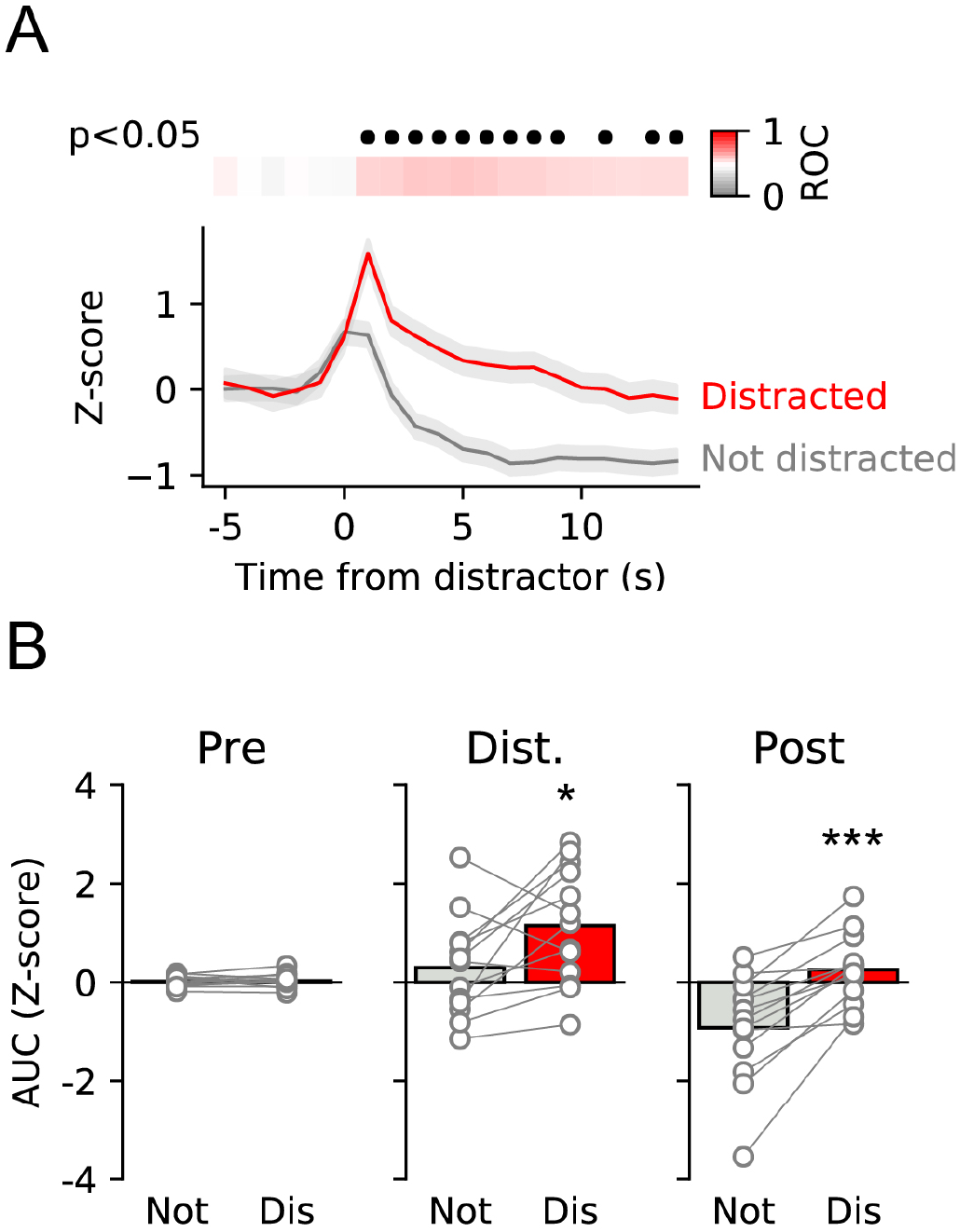
Prolonged neural activity is associated with distracted trials vs. non-distracted trials. (A) ROC analysis of photometry data from first distraction day comparing distracted and non-distracted trials shows that neural activity differs in the 15 s following presentation of a distractor. Upper panel shows ROC values coded in colour with red trials representing greater neural activity during distracted trials vs. not distracted trials. Black filled circles indicate time bins in which the ROC comparison was significant (Bonferroni corrected). (B) Comparisons of same three epochs as in Fig. 3 shows that neural activity is elevated both immediately (1-3 s; centre) and for a sustained period (4-15 s; right) following the distractor on distracted trials vs. not distracted trials. Circles show data from individual rats and bars are mean. *, *** p<0.05, <0.001 vs. not distracted trials.

In summary, neural activity in the VTA was altered not just from the presence of a distracting stimulus but also based on whether the rat paused its licking or not. However, the level of activity in VTA evoked by a distractor did not determine for how long the rat would disengage from licking behaviour.

### VTA responses to distractors do not habituate as rapidly as behaviour

Rats’ behavioural response to distracting stimuli habituated both within session and across as shown by the reduction in the likelihood of distraction and a decrease in the post-distraction pause (Fig. 1D and 1E). To test whether this change in behaviour was associated with a change in VTA neural activity we compared fibre photometry responses to the distractor between the two sessions with distractors.

When all trials were included in the ROC analysis we found no difference in the response to distractors between the distraction and habituation day in any time bin (p’s > 0.05 for all time bins; Fig. 6A). Thus, neural activity evoked by distractors does not reflect differences in behaviour observed. We proceeded to consider whether there would be differences between these two days if we separated trials into distracted or non-distracted trials. However, comparison either of only the distracted trials (Fig. 6B) or only the not distracted trials (Fig. 6C) also failed to reveal any difference between the first distraction and habituation day (for both comparisons p’s > 0.05 for all time bins). In addition, we tested whether there was evidence of within-session habituation of neural responses. However, when we compared neural responses in the three terciles from the first distraction session, we found no evidence of neural activity habituating within a session (Fig. S2; one-way repeated measures ANOVA: F(2,24)=0.78, p=0.4689).

**Figure 6.**
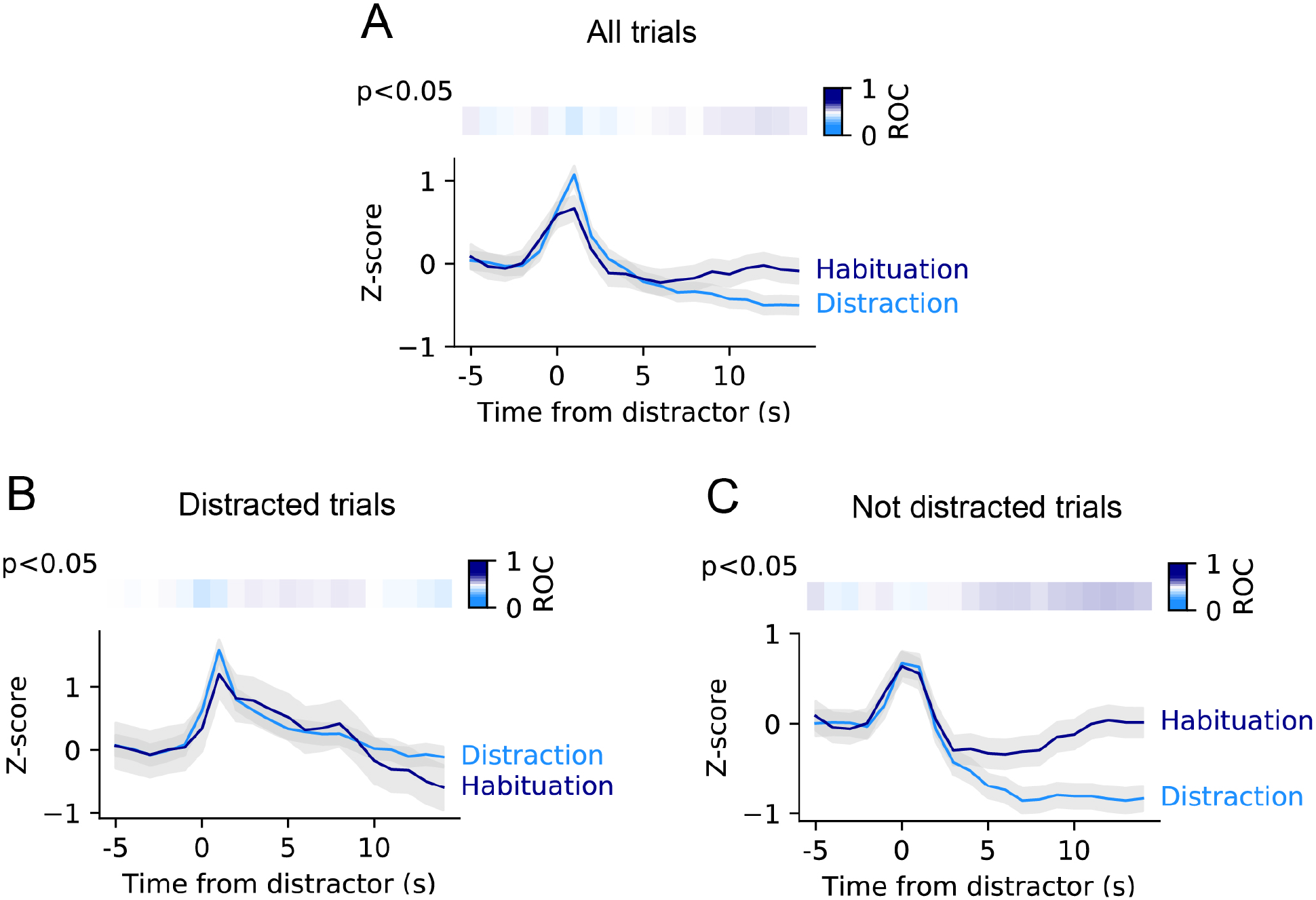
Neural activity in response to distracting stimuli does not differ between first distraction day and second distraction day when behavioural responses have habituated. (A) ROC analysis of photometry data comparing first distraction day and second distraction (habituation) day. The curves did not differ in any 1-s time bin. Upper panel shows ROC values coded in colour with dark blue bins representing greater neural activity during habituation day vs. first distraction day. The ROC comparison was not significant at any point. (B) As in A but with only trials when rat was distracted included. (C) As in A but with only trials in which the rat was not distracted included.

In a final analysis, we considered whether there were other differences in activity between the sessions that could explain the change in behaviour. One possibility is that a difference in baseline activity could be influencing the z-scored signals that are analysed and presented above. We found no difference in the RMS of the non z-scored signal between the two distraction days (Fig. S6A; paired t-test, p=0.139) but there was a trend towards a difference in the pre-distractor baseline activity of the non-z-scored signal (Fig. S6B; p=0.071). Moreover, when the post-distractor epoch of the non-z-scored signal was analysed we found a slight increase in activity evoked by the distractor on the first distraction day vs. the second day (Fig. S6C; p=0.041).

In conclusion, during ongoing licking, presentation of distracting stimuli evokes neural activity in VTA and this response does not rapidly habituate despite significant habituation of behaviour (i.e. reduced efficacy of distractor in producing a pause in licking). However, subtle changes in VTA activity in the period before the distractor might be linked to the change in behaviour.

## Discussion

Here, we have investigated how presentation of distracting stimuli during ongoing consumption affects subsequent licking behaviour and associated neural activity in VTA. In brief, we find that distracting stimuli, as predicted, cause rats to interrupt their licking and that this behavioural effect habituates such that the stimuli are less likely to distract rats on the second day they are presented. Neural activity in VTA was associated with some of these effects and was evoked by distracting stimuli. In addition, on trials in which rats were distracted, neural activity was elevated, relative to trials in which rats continued to lick. Finally, the neural signal appeared slower to habituate than the behaviour as we found no differences in the magnitude of neural activity produced on the second day distractors were presented compared to the first despite significant changes in behaviour.

Our results are broadly consistent with (O’Connor *et al*., 2015) who showed lick-triggered distracting stimuli lead to immediate interruption of licking in a large percentage of cases. Our results expand this by examining in more fine-grained detail the effect on licking behaviour and how responses to these distracting stimuli habituate over time. Here, novel stimuli (visual and auditory cues) presented during consumption lead to pauses in licking. This behaviour, and its accompanying VTA neural response, fits with salience accounts of VTA dopamine neurons which respond to neutral and novel cues. Increased VTA activity may promote exploration of potentially important stimuli in the environment by promoting orienting responses towards potential threats or opportunities. Activity of midbrain structures such as VTA is known to invigorate behaviour in reward contexts but VTA activity may also play a dual role in enabling animals to attend to stimuli in the environment to promote learning about cues and their outcomes (Nicola, 2010; Walton *et al*., 2011).

Once distractor stimuli become less novel – with repeated presentation across two sessions – they are less effective in disrupting ongoing consumption. Given that these stimuli are not predictive of either reward or punishment (they are inconsequential and the environment is stable), this is consistent with the function of dopamine neurons in reinforcement learning. The information carried by these cues is no longer driving exploration and an exploitation strategy is more fruitful, therefore the rats continue to lick for saccharin. In addition, rats may learn an association between licking and the triggering of distractor stimuli which could further decrease novelty. However, we find that the distractor-evoked VTA activity remains despite this behavioural habituation. It is possible that the VTA continues to respond to these stimuli, perhaps excited via sensory inputs from other areas (for example, via fast signals from the superior colliculus (Zhou *et al*., 2019) but that this firing response alone is not driving the disengagement from consumption. In addition, there may be subtle differences in tonic activity which are not thoroughly examined here. As such, changes to the balance between tonic and phasic activity of VTA neurons may occur and further investigation of this using other methods may yield interesting results.

A limitation of our study is that the virus we have used targets VTA neurons non-specifically. As such, we cannot ascribe these changes to a particular subset of VTA neurons. The majority of neurons in VTA are dopaminergic (Nair-Roberts *et al*., 2008; Morales & Margolis, 2017) and, in fact, it is this population of neurons that have been most extensively studied with respect to the phenomena and processes likely to be engaged by our behavioural paradigm. There is well known heterogeneity within the VTA in terms of cell type, inputs and outputs as well as functional differences in responses to stimuli (Brischoux *et al*., 2009; Lammel *et al*., 2011; Besson *et al*., 2012). As well as dopamine neurons, there is also a large population of GABA neurons some of which are interneurons and some of which are projection neurons. These too have been shown to respond to environmental stimuli and may contribute to the population response we have recorded here (Cohen *et al*., 2012). Finally, a small number of glutamate neurons are also present although their functional role remains less well defined (Hnasko *et al*., 2012; Morales & Margolis, 2017). Thus, a key question that remains how do each of these populations contribute to the population responses we show to distracting stimuli.

The VTA also receives substantial inputs from multiple areas and different transmitter systems, we do not yet fully understand how these vast inputs contribute to the modulation of VTA neurons within such tasks. In addition, cell body firing within the VTA may be separated from terminal dopamine release (Cachope *et al*., 2012; Threlfell *et al*., 2012) which is a driver of reward behaviour. The complex mechanisms of terminal modulation of dopamine release may contribute to the disparity in neural activity and behaviour which we report. Tuning down terminal dopamine release (particularly in the nucleus accumbens shell) when cues are no longer ambiguous or uncertain (i.e., when the animal has learnt they are inconsequential) may provide a mechanism for efficiently allocating attention to only the most behaviourally relevant events (Baudonnat *et al*., 2013).

Fibre photometry allows bulk population responses to be recorded. However, to simultaneously orchestrate and coordinate several phenomena the VTA may multiplex information whereby smaller subpopulations of neurons are responsible for representing and enacting different aspects of behaviour. For example, a recent paper employed single cell calcium imaging to show that VTA dopamine neurons coded sensory, motor and cognitive variables in a complex decision-making task requiring navigation (Engelhard *et al*., 2019). Dopamine release across the ventral and dorsal striatum and other projection targets may also be specific to certain phenomena with functional consequences.

A further consideration is how the internal state of the animal might influence this behaviour. We know that an animals’ energy status can substantially shape how contending behaviours are chosen, with more hungry animals more motivated to consume and less concerned with safety/threat detection (Sutton & Krashes, 2020). Future experiments could investigate how manipulations to energy status (hunger) or specific deficits (e.g. thirst, sodium or protein depletion) affect the likelihood of distractors interrupting ongoing consumption and the consequent neural activity. The design of this behavioural paradigm allows for the manipulation of consumed substances as well as fine grained analysis of licking patterns and neural activity which could begin to inform us on how different need states might modulate attention and allocation of behavioural resources.

In conclusion, our results highlight the complex role of the VTA in maintaining ongoing appetitive and consummatory behaviours while also monitoring the environment for salient stimuli. We have gone some way to examine how the VTA may encode multiple streams of information, but it remains to be determined how the VTA and other areas assimilate information about external cues with learned associations and internal states to produce adaptive behaviour to maximise positive outcomes.

## Acknowledgements

KZP was funded by the Midlands Integrated Biosciences Training Programme (BBSRC and University of Leicester). This work was supported by the Biotechnology and Biological Sciences Research Council [grant # BB/M007391/1 to JEM]; and the European Commission [grant # GA 631404 to JEM]. We would also like to acknowledge the help and support from the staff of the Division of Biomedical Services, Preclinical Research Facility, University of Leicester, for technical support and the care of experimental animals.

## Author contributions

KZP designed and conducted experiments, collected data, analysed data and wrote the manuscript, JEM designed experiments, provided data analysis scripts, wrote the manuscript and made figures. AMJY wrote the manuscript.

## Data accessibility statement

All raw data files will be published with this manuscript alongside the Python scripts used to perform analysis. These are deposited on Github (https://github.com/mccutcheonlab/distraction-peters/releases/tag/v1.0) and Figshare (https://www.doi.org/10.25392/leicester.data.12732734).

## Abbreviations

VTA: Ventral tegmental area
NAc: Nucleus accumbens
GABA: Gamma aminobutyric acid
ROC: Receiver-operator characteristic
GFP: Green fluorescent protein
PFA: Paraformaldehyde
PBS: Phosphate buffered saline

---

**Figure S1.**
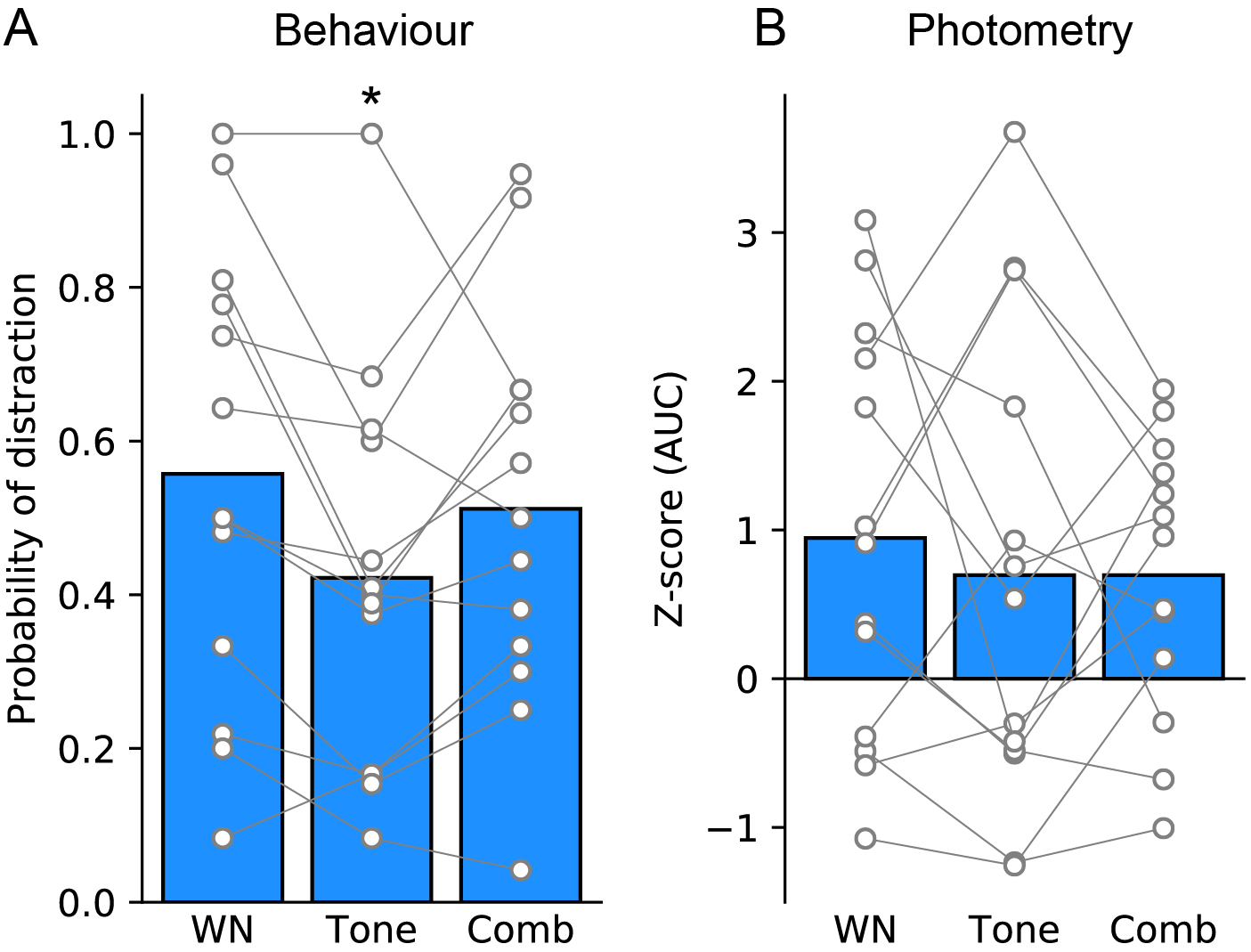
Distractors containing white noise are more distracting but neural activity is not modulated by distractor type. (A) Probability of distraction is modulated by the type of distractor. Distractors including white noise (white noise and combined) were more likely to cause distraction than distractors that just included a tone (one way ANOVA: F(2,24)=4.40, p=0.0235, post hoc Sidak’s: white noise vs tone, p=0.0256). Circles show data from individual rats for different trial types and bars are mean. (B) VTA responses were not affected by distractor type (one way ANOVA: F(2,24)=0.28, p=0.7554). Circles show data from individual rats for different trial types and bars are mean z-score AUC values taken 1-4 seconds following distractor presentation. WN, white noise; Comb, combined. *, p<0.05 vs. white noise.

**Figure S2.**
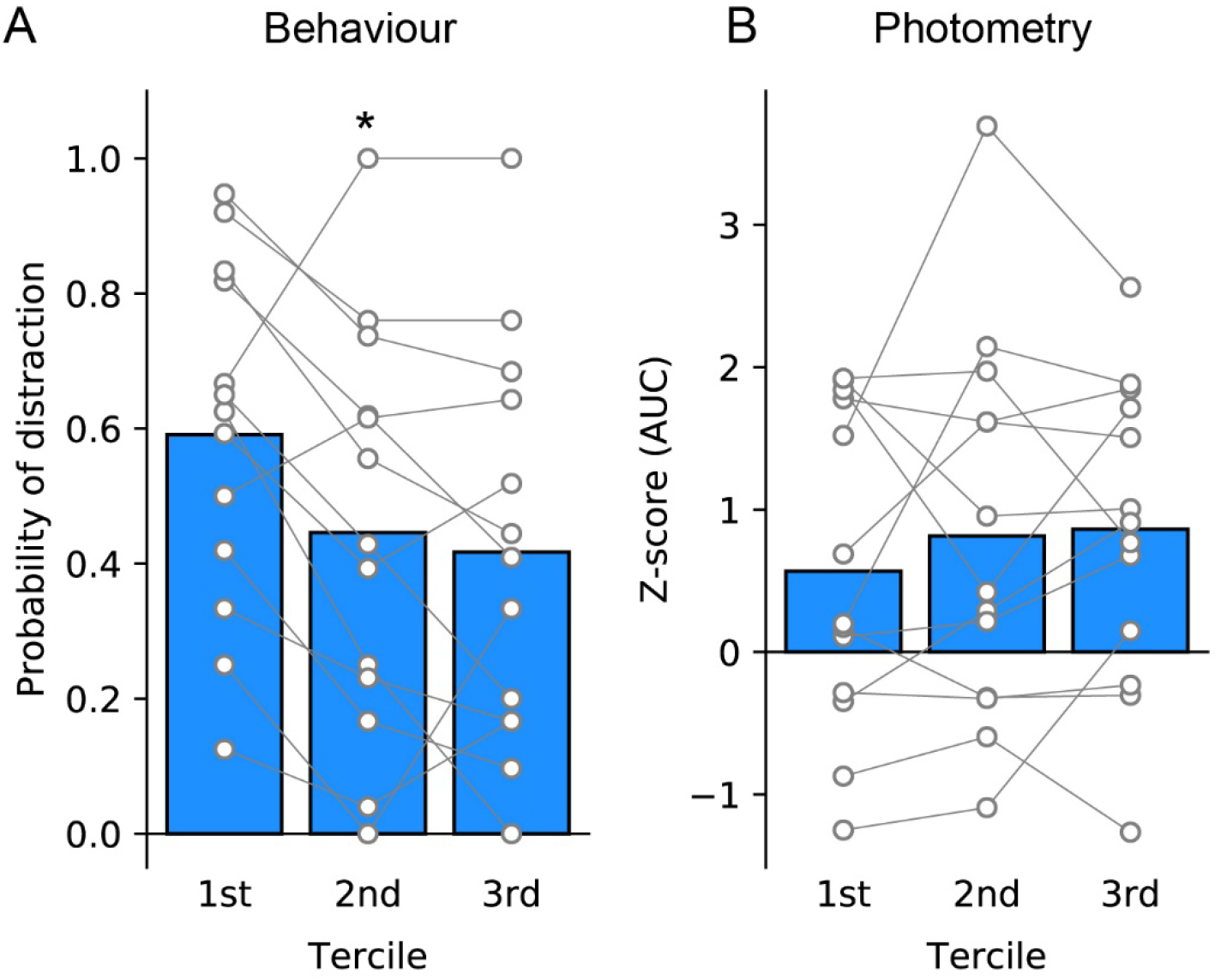
Evidence of within-session habituation. Distractors were split into terciles based on their occurrence within the distraction session (Early, 1st; Middle, 2nd; Late, 3rd). (A) Probability of distraction is reduced across the session (F(2,24)=5.01, p=0.0152). Rats were significantly more likely to be distracted in the first tercile than in the second tercile (p=0.0453). (B) Neural activity (AUC from 1-4 s following the distractor) was not different across terciles within the distraction session (F(2,24)=0.78, p=0.4689). Circles show data from individual rats in each session tercile and bars are mean. *, p<0.05 vs. 1st tercile.

**Figure S3.**
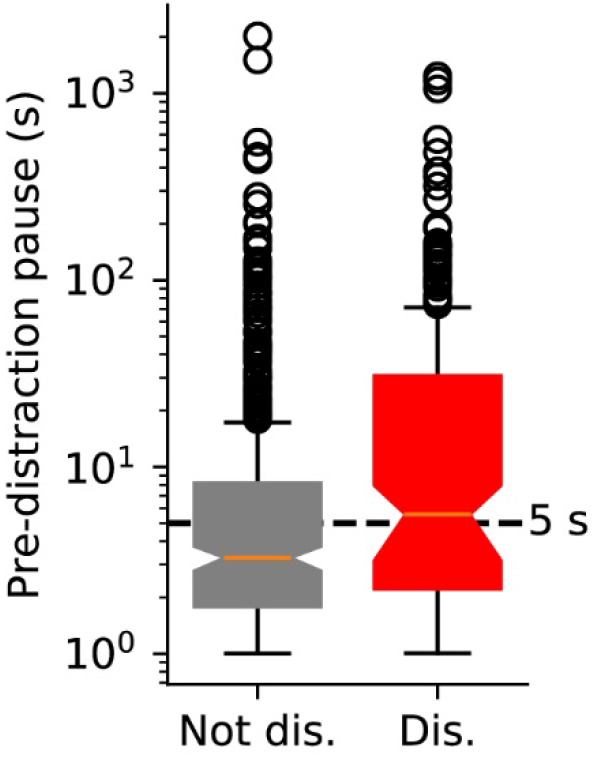
Pre-distractor pause is shorter on non-distracted trials than on distracted trials. Median pre-distractor pause was 3.27 on non-distracted trials vs. 5.57 s on distracted trials (t-test of log-transformed data, p=2×10^-7^). On 64% of non-distracted trials there was a distractor occurring in the baseline period whereas for distracted trials this proportion was 41%. Box and whisker plot shows 25-75% quartiles with whiskers at 95% quartiles. Notch and line show median and circles show data points which fall outside of the 95% confidence intervals. Dashed line shows 5s ‘baseline’ period.

**Figure S4.**
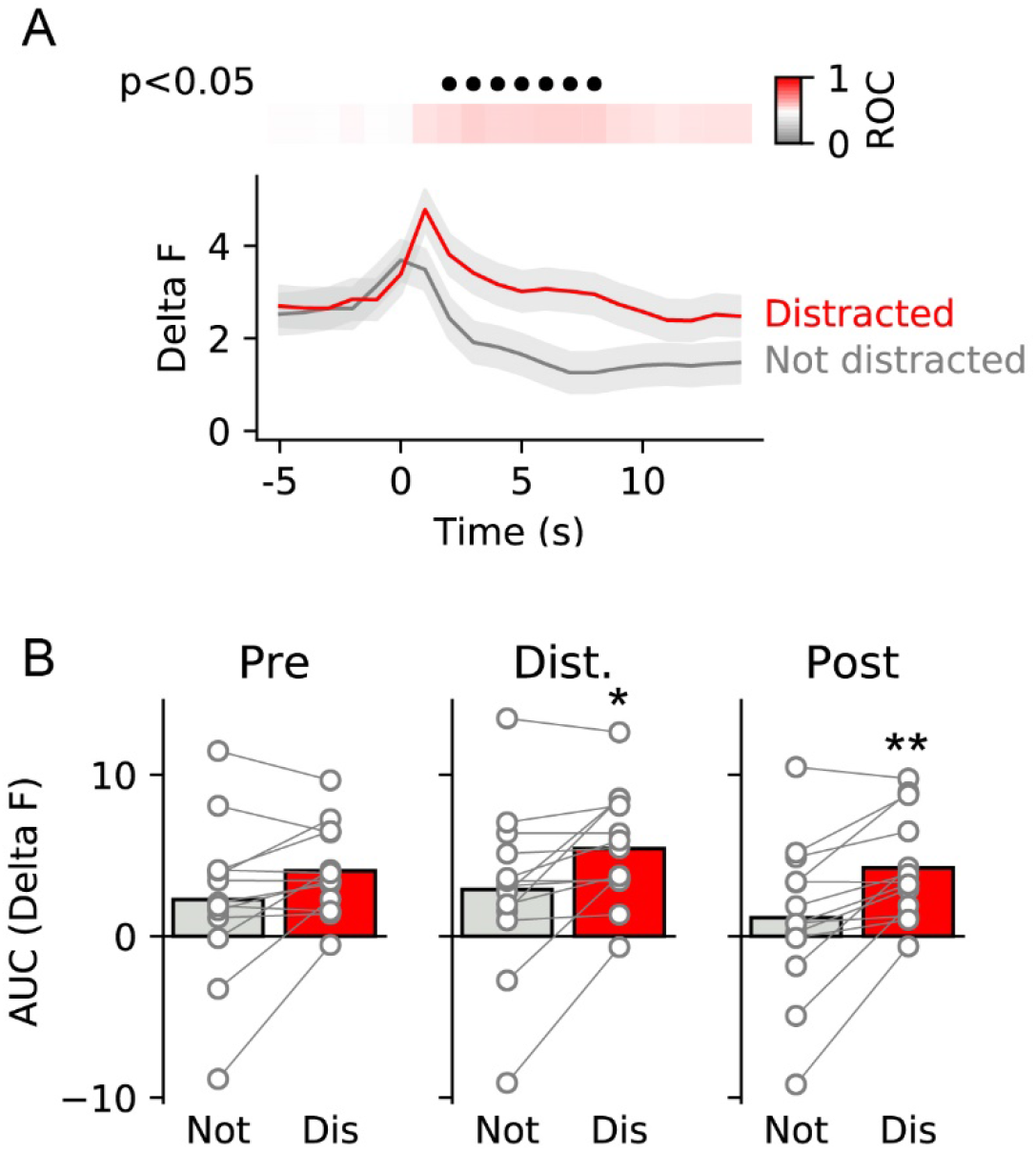
No difference in pre-di stractor neural activity on distracted and non-distracted trials. Comparison of filtered signal before z-scoring in distracted and non-distracted trials showed no difference in the epoch before the distractor (p=0.144). (A) ROC analysis of photometry data between distracted and non-distracted trials. Upper panel shows ROC values coded in colour with red trials representing greater neural activity during distracted trials vs. non-distracted trials. Black filled circ!es indicate time bins in which the ROC comparison was significant (p<0.05, Bonferroni corrected). (B) Comparisons during three epochs show there was still a difference in activity between the epochs following the distractor (immediate epoch, middle: p=0.041, late epoch, right: p=0.007) but no difference in the epoch before the distractor (pre epoch, left: p=0.144. Circles show data from individual rats and bars are mean. **, *, p<0.01, p<0.05 vs. not distracted trials.

**Figure S5.**
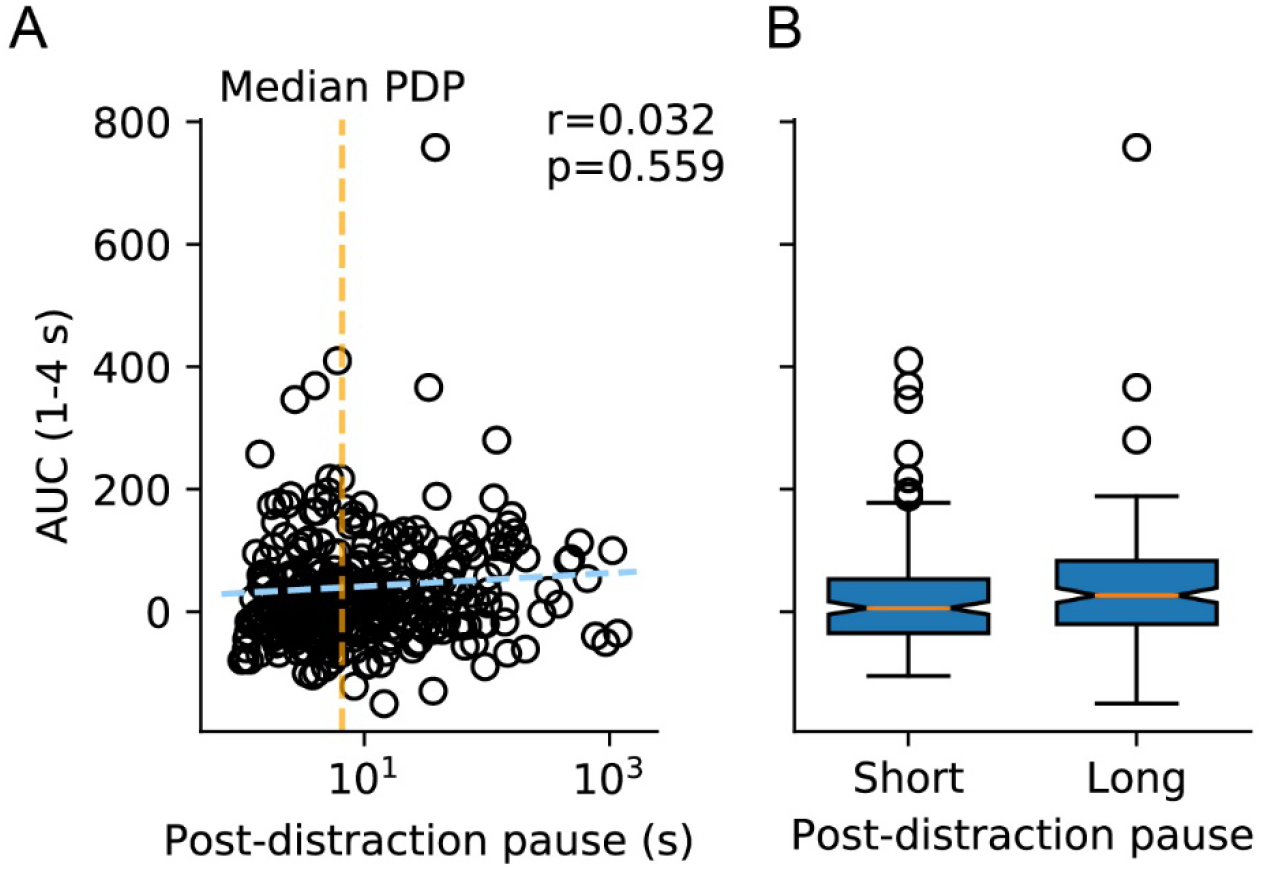
No correlation between post-distraction pause and neural activity. (A) For each trial on which rats were distracted, neural activity (AUC, 1-4 s) was regressed aginst post-distraction pause. No correlation was found. Circles show individual trials. Blue dashed line shows linear fit and orange vertical line shows median post-distraction pause (PDP). (B) When trials were median split into those with short and long post-distraction pauses, so difference in neural activity was found. Box and whisker plot shows 25-75% quartiles with whiskers at 95% quartiles. Notch and line show median and circles show data points which fall outside of the 95% confidence intervals. Dashed line shows 5s ‘baseline’ period.

**Figure S6.**
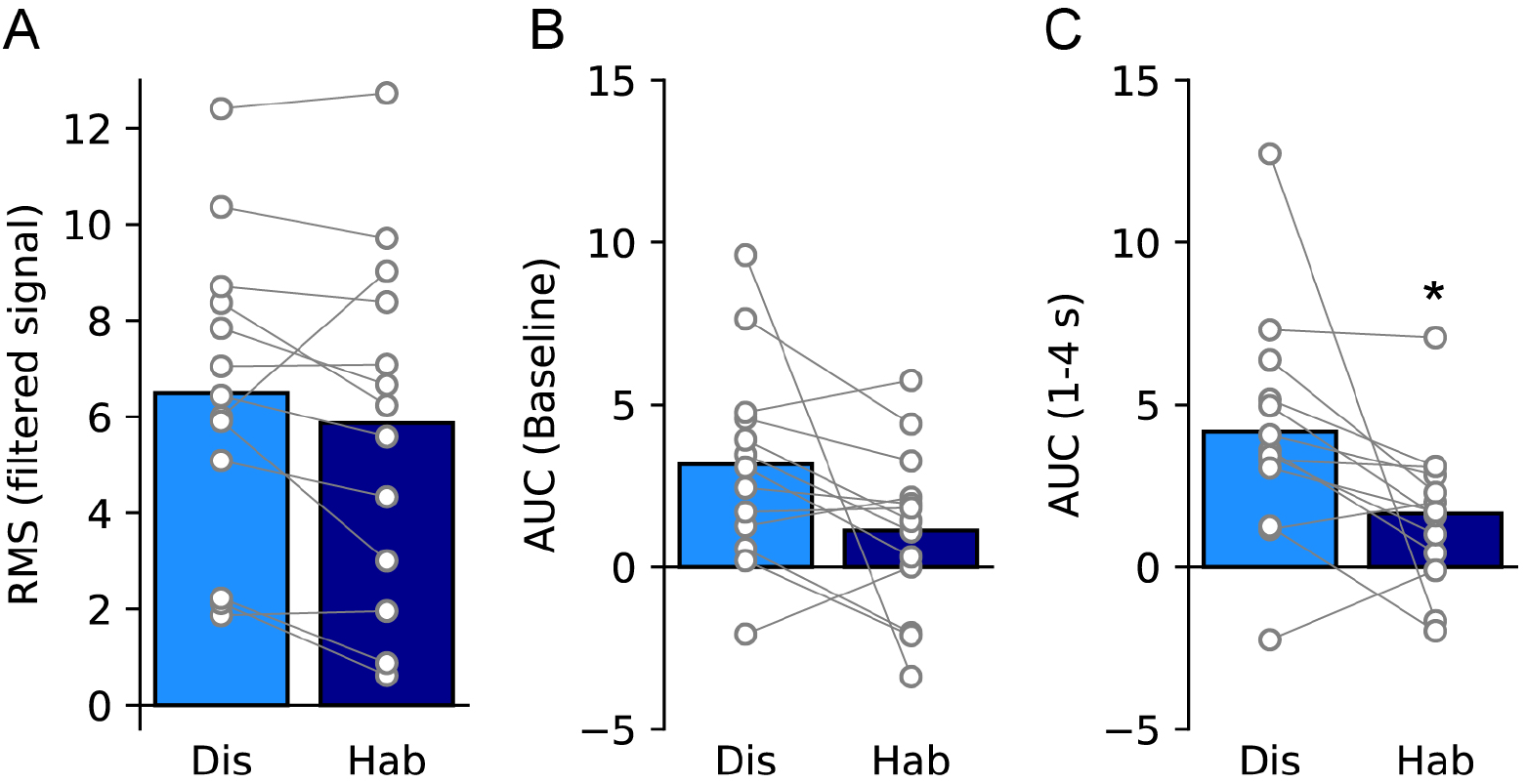
Comparison of baseline changes in photometry signal across days. (A) No difference in root mean square of filtered signal was found between the two distraction days (p=0.139). (B) There was a trend for neural activity during baseline before distractor to be higher on the first distraction day vs. the second (p=0.071). (C) When the non z-scored signal was examined, there was an elevation of activity in the immediate epoch following the distrator on the first distraction day, relative to the second (p=0.041). Circles show data from individual rats and bars are mean. *, p<0.05 vs. distraction day.

## References

Baudonnat, M., Huber, A., David, V., & Walton, M.E. (2013) Heads for learning, tails for memory: reward, reinforcement and a role of dopamine in determining behavioral relevance across multiple timescales. Front. Neurosci., 7.

Beeler, J.A., Cools, R., Luciana, M., Ostlund, S.B., & Petzinger, G. (2014) A kinder, gentler dopamine… highlighting dopamine’s role in behavioral flexibility. Front. Neurosci., 8.

Besson, M., David, V., Baudonnat, M., Cazala, P., Guilloux, J.-P., Reperant, C., Cloez-Tayarani, I., Changeux, J.-P., Gardier, A.M., & Granon, S. (2012) Alpha7-nicotinic receptors modulate nicotine-induced reinforcement and extracellular dopamine outflow in the mesolimbic system in mice. Psychopharmacology, 220, 1–14.

Brischoux, F., Chakraborty, S., Brierley, D.I., & Ungless, M.A. (2009) Phasic excitation of dopamine neurons in ventral VTA by noxious stimuli. PNAS, 106, 4894–4899.

Bromberg-Martin, E.S. & Hikosaka, O. (2009) Midbrain Dopamine Neurons Signal Preference for Advance Information about Upcoming Rewards. Neuron, 63, 119–126.

Bromberg-Martin, E.S., Matsumoto, M., & Hikosaka, O. (2010) Dopamine in motivational control: rewarding, aversive, and alerting. Neuron, 68, 815–834.

Bromberg-Martin, E.S., Matsumoto, M., Nakahara, H., & Hikosaka, O. (2010) Multiple Timescales of Memory in Lateral Habenula and Dopamine Neurons. Neuron, 67, 499–510.

Cachope, R., Mateo, Y., Mathur, B.N., Irving, J., Wang, H.-L., Morales, M., Lovinger, D.M., & Cheer, J.F. (2012) Selective activation of cholinergic interneurons enhances accumbal phasic dopamine release: setting the tone for reward processing. Cell Rep, 2, 33–41.

Cohen, J.Y., Haesler, S., Vong, L., Lowell, B.B., & Uchida, N. (2012) Neuron-type specific signals for reward and punishment in the ventral tegmental area. Nature, 482, 85–88.

Cools, R., Nakamura, K., & Daw, N.D. (2011) Serotonin and Dopamine: Unifying Affective, Activational, and Decision Functions. Neuropsychopharmacology, 36, 98–113.

Engelhard, B., Finkelstein, J., Cox, J., Fleming, W., Jang, H.J., Ornelas, S., Koay, S.A., Thiberge, S.Y., Daw, N.D., Tank, D.W., & Witten, I.B. (2019) Specialized coding of sensory, motor and cognitive variables in VTA dopamine neurons. Nature, 570, 509–513.

Fiorillo, C.D., Yun, S.R., & Song, M.R. (2013) Diversity and Homogeneity in Responses of Midbrain Dopamine Neurons. J. Neurosci., 33, 4693–4709.

Flagel, S.B., Clark, J.J., Robinson, T.E., Mayo, L., Czuj, A., Willuhn, I., Akers, C.A., Clinton, S.M., Phillips, P.E.M., & Akil, H. (2011) A selective role for dopamine in stimulus–reward learning. Nature, 469, 53–57.

Floresco, S.B. (2013) Prefrontal dopamine and behavioral flexibility: shifting from an “inverted-U” toward a family of functions. Front. Neurosci., 7.

Haluk, D.M. & Floresco, S.B. (2009) Ventral Striatal Dopamine Modulation of Different Forms of Behavioral Flexibility. Neuropsychopharmacology, 34, 2041–2052.

Hnasko, T.S., Hjelmstad, G.O., Fields, H.L., & Edwards, R.H. (2012) Ventral tegmental area glutamate neurons: electrophysiological properties and projections. J. Neurosci., 32, 15076–15085.

Horvitz, J.C., Stewart, T., & Jacobs, B.L. (1997) Burst activity of ventral tegmental dopamine neurons is elicited by sensory stimuli in the awake cat. Brain Research, 759, 251–258.

Kakade, S. & Dayan, P. (2002) Dopamine: generalization and bonuses. Neural Networks, 15, 549–559.

Lammel, S., Ion, D.I., Roeper, J., & Malenka, R.C. (2011) Projection-Specific Modulation of Dopamine Neuron Synapses by Aversive and Rewarding Stimuli. Neuron, 70, 855–862.

Lerner, T.N., Shilyansky, C., Davidson, T.J., Evans, K.E., Beier, K.T., Zalocusky, K.A., Crow, A.K., Malenka, R.C., Luo, L., Tomer, R., & Deisseroth, K. (2015) Intact-Brain Analyses Reveal Distinct Information Carried by SNc Dopamine Subcircuits. Cell, 162, 635–647.

Lisman, J.E. & Grace, A.A. (2005) The Hippocampal-VTA Loop: Controlling the Entry of Information into Long-Term Memory. Neuron, 46, 703–713.

Liu, Z., Zhou, J., Li, Y., Hu, F., Lu, Y., Ma, M., Feng, Q., Zhang, J., Wang, D., Zeng, J., Bao, J., Kim, J.-Y., Chen, Z.-F., Mestikawy, S.E., & Luo, M. (2014) Dorsal Raphe Neurons Signal Reward through 5-HT and Glutamate. Neuron, 81, 1360–1374.

Ljungberg, T., Apicella, P., & Schultz, W. (1992) Responses of monkey dopamine neurons during learning of behavioral reactions. Journal of Neurophysiology, 67, 145–163.

Moher, J., Anderson, B.A., & Song, J.-H. (2015) Dissociable Effects of Salience on Attention and Goal-Directed Action. Current Biology, 25, 2040–2046.

Morales, M. & Margolis, E.B. (2017) Ventral tegmental area: cellular heterogeneity, connectivity and behaviour. Nature Reviews Neuroscience, 18, 73–85.

Nair-Roberts, R.G., Chatelain-Badie, S.D., Benson, E., White-Cooper, H., Bolam, J.P., & Ungless, M.A. (2008) Stereological estimates of dopaminergic, GABAergic and glutamatergic neurons in the ventral tegmental area, substantia nigra and retrorubral field in the rat. Neuroscience, 152, 1024–1031.

Nicola, S.M. (2010) The flexible approach hypothesis: unification of effort and cue-responding hypotheses for the role of nucleus accumbens dopamine in the activation of reward-seeking behavior. J. Neurosci., 30, 16585–16600.

Nomoto, K., Schultz, W., Watanabe, T., & Sakagami, M. (2010) Temporally Extended Dopamine Responses to Perceptually Demanding Reward-Predictive Stimuli. J. Neurosci., 30, 10692–10702.

O’Connor, E.C., Kremer, Y., Lefort, S., Harada, M., Pascoli, V., Rohner, C., & Lüscher, C. (2015) Accumbal D1R Neurons Projecting to Lateral Hypothalamus Authorize Feeding. Neuron, 88, 553–564.

Paxinos, G. & Watson, C. (2007) The Rat Brain in Stereotaxic Coordinates, 6th edn. Elsevier.

Redgrave, P., Prescott, T.J., & Gurney, K. (1999) Is the short-latency dopamine response too short to signal reward error? Trends Neurosci., 22, 146–151.

Riccio, C.A., Reynolds, C.R., Lowe, P., & Moore, J.J. (2002) The continuous performance test: a window on the neural substrates for attention? Archives of Clinical Neuropsychology, 17, 235–272.

Salamone, J.D., Correa, M., Farrar, A.M., Nunes, E.J., & Pardo, M. (2009) Dopamine, behavioral economics, and effort. Front. Behav. Neurosci., 3.

Schultz, W. (1997) Dopamine neurons and their role in reward mechanisms. Current Opinion in Neurobiology, 7, 191–197.

Schultz, W. (2002) Getting Formal with Dopamine and Reward. Neuron, 36, 241–263.

Schultz, W. (2007) Multiple Dopamine Functions at Different Time Courses. Annual Review of Neuroscience, 30, 259–288.

Schultz, W. (2017) Reward prediction error. Current Biology, 27, R369–R371.

Steinberg, E.E., Keiflin, R., Boivin, J.R., Witten, I.B., Deisseroth, K., & Janak, P.H. (2013) A causal link between prediction errors, dopamine neurons and learning. Nature Neuroscience, 16, 966–973.

Sutton, A.K. & Krashes, M.J. (2020) Integrating Hunger with Rival Motivations. Trends in Endocrinology & Metabolism, 31, 495–507.

Threlfell, S., Lalic, T., Platt, N.J., Jennings, K.A., Deisseroth, K., & Cragg, S.J. (2012) Striatal dopamine release is triggered by synchronized activity in cholinergic interneurons. Neuron, 75, 58–64.

Waelti, P., Dickinson, A., & Schultz, W. (2001) Dopamine responses comply with basic assumptions of formal learning theory. Nature, 412, 43–48.

Walton, M.E., Gan, J.O., & Phillips, P.E.M. (2011) The influence of dopamine in generating action from motivation. In Neural Basis of Motivational and Cognitive Control. pp. 163–187.

Wang, H.-L., Zhang, S., Qi, J., Wang, H., Cachope, R., Mejias-Aponte, C.A., Gomez, J.A., Mateo-Semidey, G.E., Beaudoin, G.M.J., Paladini, C.A., Cheer, J.F., & Morales, M. (2019) Dorsal Raphe Dual Serotonin-Glutamate Neurons Drive Reward by Establishing Excitatory Synapses on VTA Mesoaccumbens Dopamine Neurons. Cell Reports, 26, 1128–1142.e7.

Winton-Brown, T.T., Fusar-Poli, P., Ungless, M.A., & Howes, O.D. (2014) Dopaminergic basis of salience dysregulation in psychosis. Trends in Neurosciences, 37, 85–94.

Wise, R.A. (2004) Dopamine, learning and motivation. Nature Reviews Neuroscience, 5, 483–494.

Zhou, Z., Liu, X., Chen, S., Zhang, Z., Liu, Y., Montardy, Q., Tang, Y., Wei, P., Liu, N., Li, L., Song, R., Lai, J., He, X., Chen, C., Bi, G., Feng, G., Xu, F., & Wang, L. (2019) A VTA GABAergic Neural Circuit Mediates Visually Evoked Innate Defensive Responses. Neuron, 103, 473–488.e6.

